# Temporal dynamics of GABA and Glx in the visual cortex

**DOI:** 10.1101/2020.01.15.907659

**Authors:** Reuben Rideaux

## Abstract

Magnetic resonance spectroscopy (MRS) can be used in vivo to quantify neurometabolite concentration and provide evidence for the involvement of different neurotransmitter systems, e.g., inhibitory and excitatory, in sensory and cognitive processes. The relatively low signal-to-noise of MRS measurements has shaped the types of questions that it has been used to address. In particular, temporal resolution is often sacrificed in MRS studies to achieve sufficient signal to produce a reliable estimate of neurometabolite concentration. Here we apply novel analyses with large datasets from human participants (both sexes) to reveal the dynamics of GABA+ and Glx in visual cortex while participants are at rest (with eyes closed) and compare this with changes in posterior cingulate cortex. We find that the dynamic concentration of GABA+ and Glx in visual cortex drifts in opposite directions, that is, GABA+ decreases while Glx increases over time. Further, we find that in visual, but not posterior cingulate cortex, the concentration of GABA+ predicts that of Glx 120 s later, such that a change in GABA+ is correlated with a subsequent opposite change in Glx. Together, these results expose novel temporal trends and interdependencies of primary neurotransmitters in visual cortex. More broadly, we demonstrate the feasibility of using MRS to investigate in vivo dynamic changes of neurometabolites.

**SIGNIFICANCE STATEMENT:** Using large datasets of magnetic resonance spectroscopy acquisitions from human participants, we develop novel analyses to investigate the temporal dynamics of neurometabolite concentration in visual cortex. We find that while participants are at rest, the concentration of GABA+ and Glx drifts in opposite directions, that is, GABA+ decreases while Glx increases over time. Further, we find that the concentration of GABA+ predicts that of Glx 120 s later, such that a change in GABA+ is correlated with a subsequent opposite change in Glx. We show these phenomena are regionally localized to visual cortex.

## INTRODUCTION

Magnetic resonance spectroscopy (MRS) can be used in vivo to measure the concentration of neurometabolites within the brain. The blood-oxygen-level-dependent signal measured using functional magnetic resonance imaging (fMRI) can provide evidence of neural activity; however, MRS can provide evidence that can be used to distinguish between different types of activity, e.g., excitatory and inhibitory. For the purpose of understanding neural mechanisms, identifying the involvement of neurotransmitter systems that support sensory/cognitive processes can be more informative than locating regions of neural representation.

Previous work using MRS to understand the role of different neurotransmitters in normal brain function has focused on neurometabolite activity in visual cortex; in particular, the function of γ-aminobutyric acid (GABA) and Glutamate (Glu), the primary inhibitory and excitatory neurotransmitters in the central nervous system, respectively. A consistent finding from these studies is that Glu increases in visual cortex in response to visual stimulation, which has been interpreted as increased involvement of this neurotransmitter during processing of visual stimuli (Bednařík et al., 2018, 2015; Ip et al., 2017; Kurcyus et al., 2018; Lin et al., 2012; Mangia et al., 2007; Schaller et al., 2013). The findings relating to GABA have been less consistent. One study found less GABA in visual cortex during visual stimulation compared to baseline (Mekle et al., 2017), while other studies have not replicated this result (Bednařík et al., 2018, 2015; Kurcyus et al., 2018; Mangia et al., 2007; Schaller et al., 2013).

The classic ‘static’ fMRS approach, i.e., comparing the average neurometabolite concentration during one experimental condition with another, has implicated GABA and Glu in visual processing. However, the information provided using this approach is severely limited. In particular, by reducing the measure of neurometabolite concentration to a single estimate averaged across a viewing condition, dynamic changes in concentration that occur under different states of visual processing are obscured. By contrast, establishing the temporal dynamics in neurometabolite concentration will reveal the timing and magnitude of change in different neurotransmitters and novel relationships between neurotransmitters. This information is essential for a comprehensive understanding of the role of neurotransmitters in visual processing, but more broadly, it may inform our understanding of the balance between excitation and inhibition, which is thought to be a key factor in multiple neurological and psychiatric illnesses (Bradford, 1995; Kehrer et al., 2008; Rubenstein and Merzenich, 2003).

The temporal resolution of MRS is highly restrained by the signal-to-noise ratio of measurements acquired using the technique. Although the duration that restricts the temporal resolution of MRS (i.e., the relaxation time) can be similar to that of fMRI (i.e., ~2 s), in order to yield a reliable measurement of neurometabolite concentration from within the brain, multiple transients must be combined to reach a sufficient signal-to-noise ratio (Mikkelsen et al., 2018). For example, it is common to combine between 200-300 transients (~10 min) to produce a single measure of neurometabolite concentration (Kurcyus et al., 2018; Rideaux and Welchman, 2018). Here we overcome the signal-to-noise limitation of MRS by applying temporal analyses of neurometabolite concentration to a large dataset of participants. We measure the dynamic concentration of GABA and Glx in visual cortex of participants while at rest (with closed eyes). We assess the regional specificity of temporal dynamics by comparing these results with data from posterior cingulate cortex (Mikkelsen et al., 2019, 2017). To observe the temporal dynamics of neurometabolites, we analyse the data in two ways. We first take a moving average of a ~6 min period, to reveal low frequency trends in the data (Chen et al., 2017; Rideaux et al., 2019). Next, using a new technique, we combine data across participants (rather than time), which allows us to track the concentration of neurometabolites with relatively high temporal resolution (12 sec) over a 13 min period.

Based on previous empirical evidence, we may expect Glx to decrease and GABA to increase under conditions of no visual stimulation. By contrast, our analyses reveal the opposite pattern of results: Glx increases while GABA decreases (in visual, but not posterior cingulate cortex). These results are broadly consistent with findings from visual deprivation studies (Boroojerdi et al., 2000; Lunghi et al., 2015) and may provide a link between the conflicting results from MRS studies that use relatively short periods of visual deprivation compared to those with longer periods. Further, we expose large changes in GABA and Glx, previously obscured by averaging over long durations and reveal a striking relationship between GABA and Glx in visual cortex: a change in GABA predicts the opposite change in Glx ~120 sec later.

## METHODS

### Participants

Fifty-eight healthy participants with normal or corrected-to-normal vision participated in the experiment. The mean age was 24.4 yr (range, 19.4–40.5 yr; 31 women). Participants were screened for contra-indications to MRI prior to the experiment. All experiments were conducted in accordance with the ethical guidelines of the Declaration of Helsinki and were approved by the University ethics committee and all participants provided informed consent.

### Data collection

Participants underwent a MR spectroscopic acquisition targeting visual cortex. During the acquisition, the lights in the room were turned off and participants were instructed to close their eyes. To compare these data with those measured from another brain region, we reanalysed previously gathered MR spectroscopic data targeting posterior cingulate cortex (Mikkelsen et al., 2019, 2017).

### Data acquisition

Magnetic resonance scanning targeting visual cortex was conducted on a 3T Siemens Prisma equipped with a 32-channel head coil. Anatomical T1-weighted images were acquired for spectroscopic voxel placement with an ‘MP-RAGE’ sequence. For detection of GABA+ and Glx, spectra were acquired using a MEGA-PRESS sequence (Mescher et al., 1998, 1996): TE=68 ms, TR=3000 ms; 256 transients of 2048 data points were acquired in 13 min experiment time; a 14.28 ms Gaussian editing pulse was applied at 1.9 (ON) and 7.5 (OFF) ppm. Water suppression was achieved using variable power with optimized relaxation delays (VAPOR; Tkáč and Gruetter, 2005) and outer volume suppression. Automated shimming followed by manual shimming was conducted to achieve approximately 12 Hz water linewidth.

The “Big GABA” dataset comprises a collection of MRS datasets collected by different groups using the same parameters on GE, Phillips, and Siemens scanners. This dataset was acquired separately for a previous study and targeted posterior cingulate cortex; thus, here it was reused as a region specificity control for the main visual cortex data. A detailed description of the data acquisitions for the Big GABA dataset can be found in Mikkelsen et al. (2019, 2017). To summarize the procedure, magnetic resonance scanning targeting posterior cingulate cortex was conducted on 3T Siemens, GE, and Phillips scanners equipped with 8-, 32-, or 64-channel head coils. Spectra were acquired using a MEGA-PRESS sequence: TE=68 ms, TR=2000 ms; 320 transients of either 2048 or 4096 data points were acquired in 12 min experiment time; a 15 ms Gaussian editing pulse was applied at 1.9 (ON) and 7.5 (OFF) ppm. The Big GABA dataset comprises sub-datasets collected by different research groups at different facilities. The following sub-datasets from the Big GABA dataset were used in the current study: G4, G5, G7, G8, P1, P3, P4, P5, P6, P7, P8, P9, P10, S1, S6, & S8; the letter in the notion refers to the scanner make and the number refers to the group that collected the data. Each sub-dataset (e.g., G4) comprises data from between 8-12 participants; in total there were data from 196 participants. With the exception of G1 and G6, this includes all the publicly available Big GABA datasets. We used the macromolecule unsuppressed transients from these datasets for closer comparison with visual cortex data, and we excluded data from G1 and G6 as these had fewer transients.

Spectra were acquired from a location targeting visual, i.e., V1/V2 (**Fig. 1a**), and posterior cingulate cortices (**Fig. 1b**); note, the posterior cingulate cortex data were acquired from the (previously described) publicly available Big GABA dataset. The voxel targeting visual cortex (3×3×2 cm) was placed medially in the occipital lobe; the lower face aligned with the cerebellar tentorium and positioned so to avoid including the sagittal sinus and to ensure it remained within the occipital lobe. The voxel targeting posterior cingulate cortex (3×3×3 cm) was positioned in the medial parietal lobe and rotated in the sagittal plane to align with a line connecting the genu and splenium of the corpus callosum. The coordinates of the voxel location were used to draw a mask on the anatomical T1-weighted image to calculate the volume of grey matter, white matter, and cerebrospinal fluid within each voxel. Segmentation was performed using the Statistical Parametric Mapping toolbox for MATLAB (SPM12, http://www.fil.ion.ucl.ac.uk/spm/).

**Figure 1.**
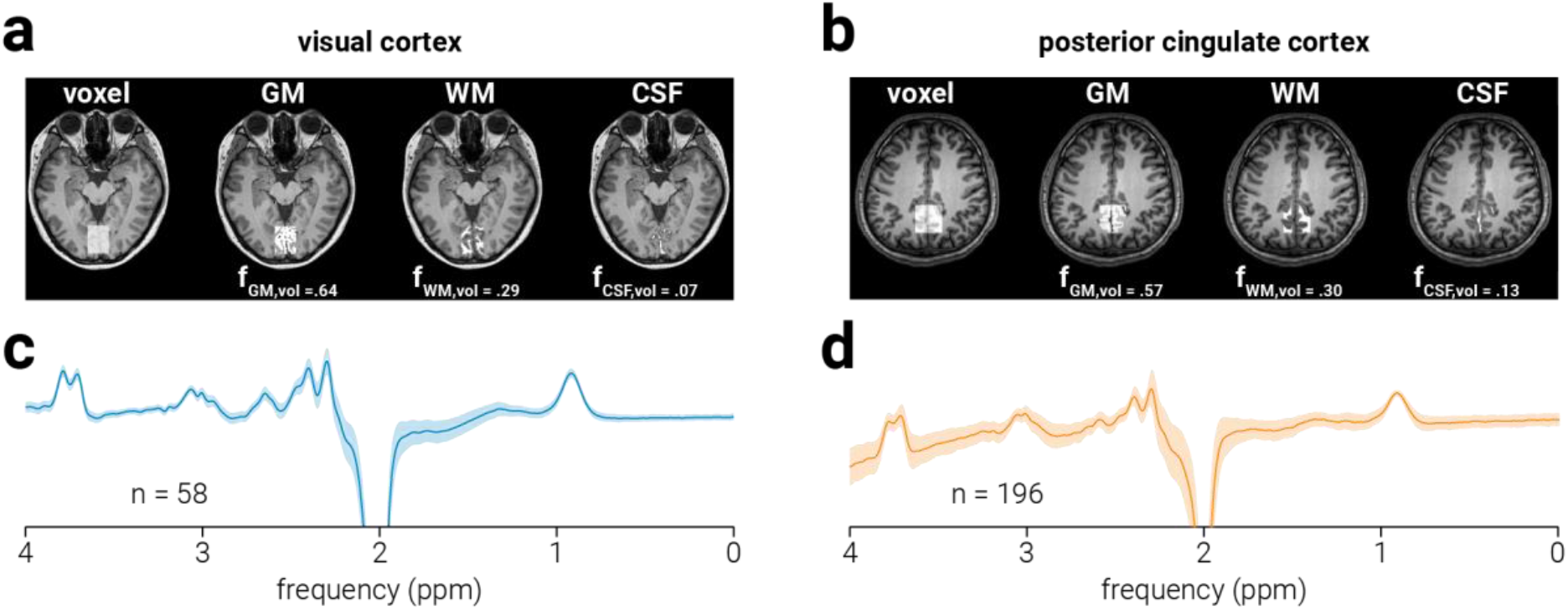
Data acquisition. Representative MRS voxel placement for **a**) visual and **b**) posterior cingulate cortices on a T1-weighted structural image and probabilistic partial volume voxel maps following tissue segmentation for representative participant. Corresponding tissue proportions of grey matter (GM), white matter (WM) and cerebrospinal fluid (CSF) are shown. Average spectra across all subjects for **c**) visual and **d**) posterior cingulate cortices; number of subjects comprising each average spectrum is shown and grey shaded regions indicate standard deviation. Note the non-uniform baseline in (**d**), this resulted from the incomplete removal of the water peak (which was supressed in the visual but not the posterior cingulate cortex data) in the difference spectra; importantly, this non uniformity was modelled and remove during neurometabolite quantification.

### Data processing

Spectral quantification was conducted in MATLAB using GANNET v3.1 (Edden et al., 2014) and in-house scripts. Frequency, phase, area, and full width at half maximum (FWHM) parameters of the Creatine peak at 3.0 ppm were estimated by fitting a Lorentzian peak to the data and individual spectra with parameter estimates >3 standard deviations from the mean were omitted from further analysis; the remaining spectra were frequency and phase corrected using these parameters. To ensure spectral alignment between transients across time and participants, spectra were aligned such that the Cr peak was centered at the same frequency (3.0 ppm). Total creatine (tCr) and total N-acetylaspartate (tNAA) signal intensity were determined by fitting a single Lorentzian peak to the mean OFF spectra at 3.0 ppm and 2.0 ppm, respectively. ON and OFF spectra were subtracted to produce the edited spectrum (**Fig. 1c** & **d**), from which GABA+ (GABA and co-edited macromolecules; 3 ppm) and Glx (a complex comprising Glu and Glutamine (Gln); 3.8 ppm) signal intensity were modelled off single- and double-Gaussian peaks, respectively. All neurometabolite signal intensities were calculated as the area of the fitted peak/s.

Data in which the FWHM of any of the quantified neurometabolites (tCr, tNAA, GABA+, or Glx) was >3 standard deviations from the mean across each dataset (e.g., visual/posterior cingulate cortex) were omitted from further analysis. This resulted in omission of data from one participant in the visual cortex dataset and three participants in the posterior cingulate cortex dataset.

Intensities of GABA+, Glx, and tNAA were normalized to the commonly used internal reference tCr (Jansen et al., 2006), yielding relative concentration values (i.e., GABA+:tCr, Glx:tCr, and tNAA:tCr; **Fig. 1e**). The tCr signal is acquired within the same MEGA-PRESS transients as the target neurometabolites. Thus, normalization of GABA+, Glx, and tNAA to tCr minimizes the influence of changes that occur during the acquisition which alter the entire spectrum, e.g., changes in signal strength, line width, chemical shift displacement, or dilution associated with changes in blood flow (Ip et al., 2017). For correlational analyses reported in **Table 2** and static analyses shown in **Figure 2**, a cerebrospinal fluid tissue-correction (Harris et al., 2015) was applied to the neurometabolite measurements with the following equation:

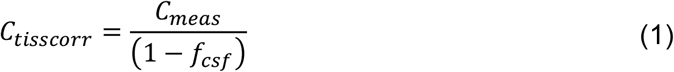

where *C_tisscorr_* and *C_meas_* are the tissue-corrected and uncorrected neurometabolite concentrations (e.g., GABA+:tCr), respectively, and *f_csf_* is the proportion of cerebrospinal fluid within the voxel. All other analyses report concentrations as a proportion of their initial magnitude, and thus do not require tissue correction.

**Table 1.** Measures of spectral quality and fit error.

| Location | FWHM (Hz) |  |  |  | Frequency drift<br>(ppm std) | Fit error (percent) |  |  |  |
| --- | --- | --- | --- | --- | --- | --- | --- | --- | --- |
|  | GABA+ | Glx | tNAA | tCr |  | GABA+ | Glx | tNAA | tCr |
| VC | 24.9±1.7 | 17.8±0.4 | 8.4±0.9 | 8.5±0.7 | 0.0088±0.0044 | 7.6±2.1 | 1.3±0.3 | 3.3±0.3 | 3.6±0.2 |
| PCC | 20.5±1.7 | 16.8±0.7 | 8.2±0.8 | 7.6±0.7 | 0.0065±0.0061 | 6.9±1.7 | 2.1±1.5 | 3.1±1.3 | 4.5±0.5 |

Note: values indicate across subject averages ± standard deviation.

**Figure 2.**
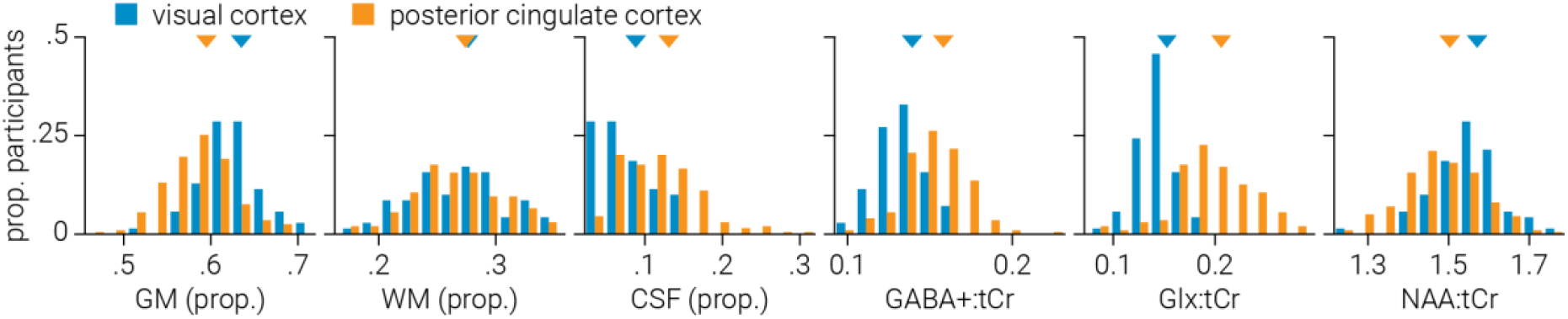
Static analysis. Distribution of grey matter (GM), white matter (WM), and cerebrospinal fluid (CSF) voxel tissue proportions, and (GABA+, Glx, tNAA) neurometabolite concentrations across participants for visual and posterior cingulate cortices. Neurometabolite concentrations are tissue-corrected and expressed as a ratio to tCr. Triangles indicate distribution means.

### Low-resolution dynamic analysis

For the low-resolution dynamic analysis of the visual cortex data, we used a sliding window (width, 128 transients; step size, 2 transients) to measure average neurometabolite concentration as it changed over the course of the scan (256 transients/768 sec). This window size was based on previous work using this analysis on similar data (Rideaux et al., 2019). For the posterior cingulate cortex data (320 transients/640 sec), we matched the duration of the sliding window width to the that used for the visual cortex data by including more transients (width, 192 transients; step size, 2 transients).

To determine if the average change in neurometabolite concentration was significantly different from zero, a two-sided *t*-test (alpha, .05) was performed at each time point. That is, for each brain region, each *t*-test included a single estimate from each participant, which corresponded to the participant’s change in neurometabolite concentration at that time. As multiple (65) tests were conducted for each neurometabolite, to reduce the likelihood of spurious significant differences in the time course, a cluster correction was applied at the group level, where time was the clustered dimension. Clusters were defined by the sum of their constituent *t*-values and compared to a null hypothesis distribution of clusters produced by shuffling the time labels (5000 permutations); positive and negative *t*-value clusters were treated separately. Clusters below the 95_th_ percentile of the null hypothesis distribution were disregarded.

### High-resolution dynamic analysis

The temporal resolution of MRS is severely limited by the signal-to-noise ratio of individual spectra. That is, to achieve the signal-to-noise ratio required to yield an accurate neurometabolite measurement, many individual spectra must be combined. In order to achieve sufficiently high signal in the low-resolution dynamic analysis, we combined multiple (128) spectra within the same subject. This method produces a dynamic trace for each participant; however, the smoothing produced by the sliding window approach may obscure both the true magnitude of metabolic change over time and dynamic changes occurring at higher frequencies. Thus, to achieve higher temporal resolution, we averaged individual ON and OFF spectra across participants to produce a single trace, with no smoothing, for each condition. As in the low temporal resolution analysis, we matched the temporal resolution between datasets in the high temporal resolution analysis (12 s) by using a resolution of four and six transients in the visual and posterior cingulate cortices, respectively.

To test for predictive relationships between GABA+ and Glx, we ran a cross-correlation analysis between the abovementioned high-resolution neurometabolite traces. We tested for relationships in both directions, that is, whether GABA+ concentration predicts Glx concentration and vice versa. The neurometabolite traces comprise a limited number of time points (visual cortex: 64, posterior cingulate cortex: 53) and there is an inverse relationship between the lag separating the neurometabolites and the number of time points included in the cross correlation analysis. This results in less reliable correlation values at lags close to the maximum duration of the neurometabolite traces due to insufficient sample sizes. To avoid these unreliable correlations, we only included lags with a minimum of 25 time points in the analysis (Bonett and Wright, 2000). This yielded a total of 40 correlations for each predictive direction between GABA+ and Glx in visual cortex and 29 for posterior cingulate cortex. Given that multiple tests were conducted, to reduce the likelihood of spurious significant correlation values in the cross-correlation analyses, a cluster correction was applied. Clusters were defined by the sum of their constituent *t*-values and compared to a null hypothesis distribution of clusters produced by shuffling the time labels (5000 permutations); positive and negative *t*-value clusters were treated separately. Clusters below the 95_th_ percentile of the null hypothesis distribution were disregarded.

### Significance testing

The significance of differences between data from different voxel locations was assessed using the independent samples *t*-test, and the significance of changes in neurometabolite concentration (from zero change) was assessed using the one-sample *t*-test; all tests were two-sided and used an alpha = .05. The normality assumption was tested with the Shapiro–Wilk test of normality. For data in which the assumption of normality was violated, significance was assessed using the Wilcoxon rank sum test. The significance of correlations between neurometabolite concentrations was assessed using the Pearson linear correlation and the Pearson linear partial correlation; all tests used an alpha = .05.

## RESULTS

### Spectra quality

**Table 1** shows the average FWHM, frequency drift and fit error for measurements taken from visual and posterior cingulate cortices. For each subject, the FWHM was calculated from the average spectra, i.e., spectra averaged across all (256/320) transients. Frequency drift was calculated as the standard deviation of the position of the Cr peak across individual OFF spectra, prior to alignment; the frequency drift values shown in **Table 1** reflect the average standard deviation across participants. The fit error for GABA+, Glx, tNAA, and tCr were divided by the amplitude of their fitted peaks to produce normalized measures of uncertainty. The average fit error for each neurometabolite was relatively low (Mullins et al., 2014; **Table 1**). Comparison between data from visual and posterior cingulate cortices revealed higher FWHM estimates for all quantified neurometabolites in visual cortex (GABA+, *t*_(248)_=16.80, *P*=8.4e_−43;_ Glx, *z*=9.24, *P*=9.4e_−20_; tNAA, *z*=2.06, *P*=.040; tCr, *z*=7.07, *P*=1.5e_−12_). This suggests that shimming may have been more effective in the posterior cingulate dataset; however, differences in voxel size, voxel tissue composition, and the number of transients collected may have contributed to this difference. We also found frequency drift (*z*=5.16, *P*=2.5e_−7_), and fit error for GABA+ (*z*= 2.59, *P*=.010) and tNAA (*z*=4.89, *P*=1.0e_−6_) were higher in the visual cortex dataset. By comparison, fit error was lower in visual cortex for Glx (*z*=-5.15, *P*=2.6e_−7_) and tCr (*z*=-10.92, *P*=9.3e_−28_).

### Static analysis

**Figure 2** shows the distribution of grey matter, white matter, and cerebrospinal fluid voxel composition across participants for the visual and posterior cingulate cortex datasets. We found that visual cortex voxels composed less grey matter (*t*_(248)_=7.53, *P*=9.3e_−13_) and more cerebrospinal fluid (*z*=-6.27, *P*=3.6e_−10_) than those from posterior cingulate cortex; no significant difference was found between white matter (*t*_(248)_=0.27, *P*=.787). We quantified the concentration of GABA+, Glx, and tNAA using the classic ‘static’ approach, in which all transients are averaged together to extract a single estimate. After applying a cerebrospinal fluid tissue-correction, we compared the average neurometabolite concentration across participants between visual and posterior cingulate cortices. We found lower concentrations of GABA+:tCr (*t*_(248)_=-6.53, *P*=3.7e_−10_) and Glx:tCr (*z*=-9.33, *P*=1.0e_−20_) in visual cortex, but higher concentration of tNAA:tCr (*t*_(248)_=4.62, *P*=6.1e_−6_; **Fig. 2**).

### Low-resolution temporal dynamics of GABA and Glx

Using a sliding temporal window analysis, we quantified change in the concentration of GABA+ and Glx measured from MRS voxels targeting visual and posterior cingulate cortices over the course of 13 and 12 min periods, respectively. We found that in visual cortex, GABA+ significantly decreased (max difference=-5.0%, *t*_(56)_=-3.25, *P*=.002) while Glx significantly increased (max difference=2.7%, *t*_(56)_=3.74, *P*=4.4e_−4_) over the course of the period (**Fig. 3a**). By comparison, we found that in posterior cingulate cortex there was no significant change in either GABA or Glx (**Fig. 3b**). The FWHM of the neurometabolite peaks indicate spectra from posterior cingulate cortex had better signal quality; further, this dataset contained more than three times as many participants as the visual cortex dataset. Thus, it seems unlikely that the neurometabolite drift we observed in visual cortex was also present in posterior cingulate cortex, but that we failed to detect it because of poor signal quality or lack of statistical power. A more parsimonious explanation is that the neurometabolite drift in visual cortex was regionally specific. Given that we referenced GABA+ and Glx to tCr, a possible concern is that the changes in neurometabolite concentration observed in visual cortex reflect changes in tCr, as opposed to GABA+ or Glx. However, this is unlikely as we did not find any change in tNAA concentration, for either visual or posterior cingulate cortices, which was also referenced for tCr (**Fig. 2a**).

**Figure 3.**
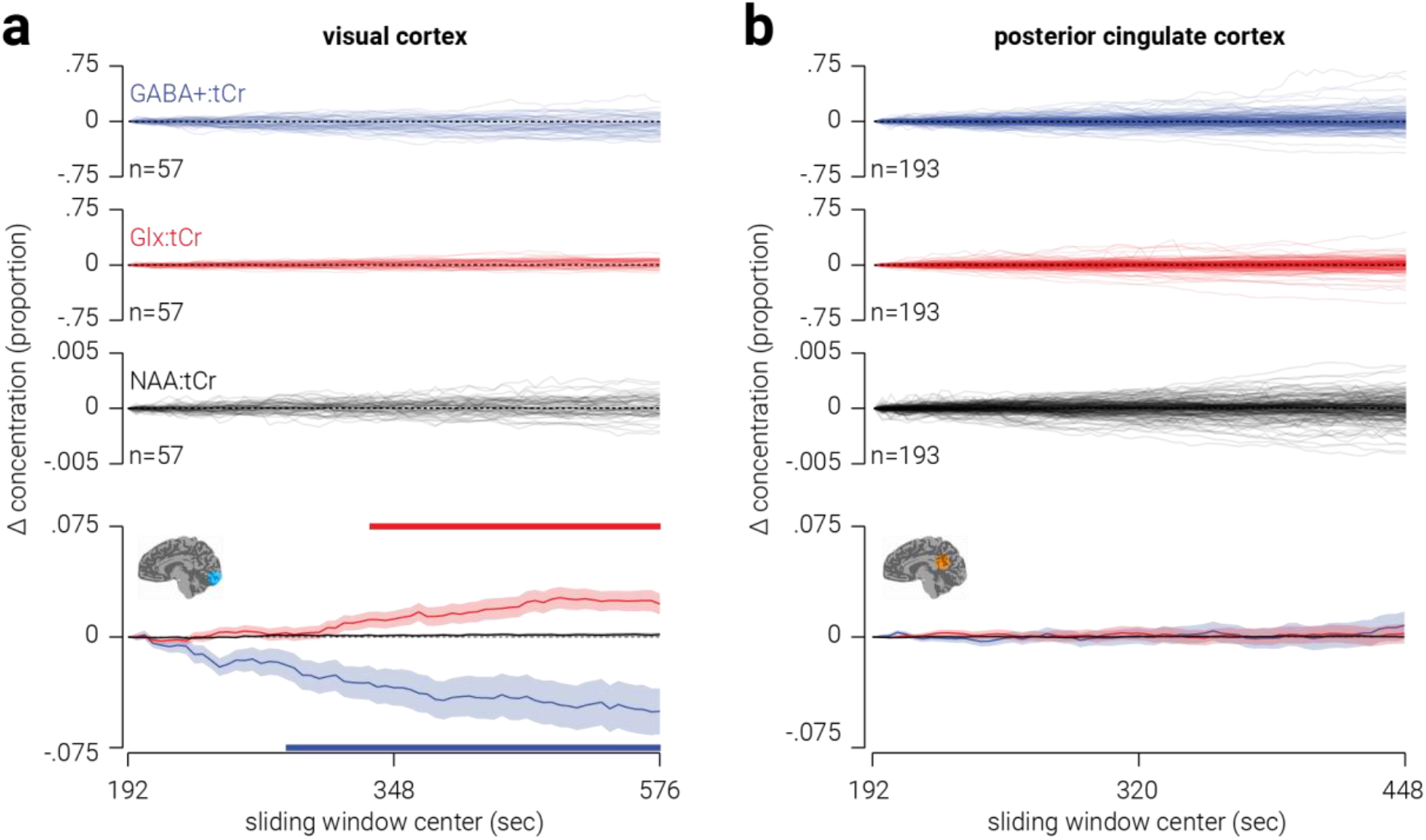
Low-resolution temporal dynamics of neurometabolites in visual and posterior cingulate cortices. (**a**, top) Individual traces showing change in GABA+, Glx, and tNAA (all referenced to tCr) measured from an MRS voxel targeting visual cortex. (**a**, bottom) Same as (**a**, top), but averaged across participants. (**b**) Same as (**a**), but from a voxel targeting posterior cingulate cortex. Shaded regions indicate s.e.m. and horizontal coloured bars at the top and bottom of (**a**) indicate (cluster corrected) periods of significant difference from zero. Note, the scale of change was smaller for tNAA:tCr than the other two neurometabolites, so it is closely overlaid with the dashed ‘zero’ line.

We next assessed whether the fit error of the GABA+, Glx, tNAA, and tCr peaks changed over the course of the acquisition. For visual cortex, the were no significant changes in the fit error of the tNAA or tCr peaks. However, there were significant increases in fit error for GABA+ between 360-394 s, and for Glx between 564-576 s. A possible concern is that the changes we observed in the concentration of GABA+ and Glx in visual cortex were due to reduced neurometabolite quantification accuracy, i.e., increased fit error. To test this possibility, we assessed the inter-individual relationship between change in fit error and change in neurometabolite concentration, for GABA+ and Glx in visual cortex. We found no evidence for a relationship between fit error and concentration for either neurometabolite (GABA+, n=57, Pearson *r*=-.14, *P*=.308; Glx, n=57, Pearson *r*=.06, *P*=.676), indicating that change in quantification accuracy does not explain for the neurometabolite drift we observed in visual cortex. For posterior cingulate cortex, we found no dynamic change in fit error for any of the neurometabolites, with the exception of the GABA+ peak, where there was a window between 196-208 s in which the fit error was significantly reduced.

Another possible concern is that changes observed in the difference spectra over time are related to scanner field drift due to gradient cooling (Lange et al., 2011) or participant motion (Bhattacharyya et al., 2007). In particular, if the scanner field drifts, the position of the editing pulse relative to the GABA and Glx peaks changes. This may change the efficiency with which the peaks are edited and thus their magnitude in the difference spectrum. As the frequency drift of the visual cortex spectra was higher than in posterior cingulate cortex, where we did not observe any change in neurometabolite concentration, this may account for the changes in neurometabolite concentration observed in visual cortex. However, if the scanner field drifted, the position of the editing pulse (1.9 ppm) relative to the GABA (3.0 ppm) and Glx (3.8 ppm) peaks would shift in the same direction for both neurometabolites. This would produce either a reduction or increase in the magnitude of both peaks. Thus, as we found GABA and Glx concentration changes in opposite directions over time, this cannot be explained by a drift in the scanner field. It is possible that changes in one of the neurometabolites could be accounted for by scanner field drift. For example, if resting induced a large decrease in GABA and no change in Glx, then scanner drift produced an increase in both peaks, if the reduction in GABA was larger than the subsequent increase due to scanner drift, the final outcome would be an increase in Glx and an attenuated decrease in GABA. As a test of this possibility, we measured the tNAA trough in the difference spectra using an inverse Lorentzian. Like the magnitude of the GABA+ and Glx peaks, the magnitude of the tNAA trough reflects the efficiency of the editing pulse. Thus, if scanner drift is responsible for changes in the magnitude of the GABA+ or Glx peaks, we would expect to see corresponding changes in the amplitude of the edited tNAA trough. However, we found no evidence for change in the amplitude of the edited tNAA trough over time. As a further test of this possibility, we assessed whether there was an inter-individual relationship between the degree of frequency drift and change in either GABA+ or Glx. If scanner drift produced the change in neurometabolite concentration we observed in visual cortex, we would expect these measures to be positively correlated. By contrast, we found no relationship between frequency drift and either change in GABA+ (n=57, Pearson *r*=-.10, *P*=.457) or Glx (n=57, Pearson *r*=.06, *P*=.662). These results indicate that scanner drift did not contribute to the changes in neurometabolite concentration.

### Correlational analyses of static and low temporal resolution measurements

For each voxel location, we assessed whether there were inter-individual relationships between different static neurometabolite concentrations or changes in concentration. For static measurements, we averaged across all 256/320 transients. The results of the analysis are shown in **Table 2**. For static neurometabolite concentrations, we found a positive relationship between Glx and tNAA measured from visual and posterior cingulate cortices. We also found a positive correlation between GABA+ and tNAA in visual cortex, and between GABA+ and Glx in posterior cingulate cortex. By contrast, the only relationship between change in metabolite concentration found was for GABA+ and tNAA in posterior cingulate cortex.

**Table 2.** Correlation coefficients between neurometabolites measured from visual and posterior cingulate cortices.

| Location | GABA+ & Glx | GABA+ & tNAA | Glx & tNAA | ΔGABA+ &<br>ΔGlx | ΔGABA+ &<br>ΔtNAA | ΔGlx &<br>ΔtNAA |
| --- | --- | --- | --- | --- | --- | --- |
| VC | -0.16 | 0.33* | 0.33* | 0.06 | 0.10 | -0.17 |
| PCC | 0.25*** | 0.50*** | 0.17* | 0.03 | -0.18* | 0.04 |

*Note: Correlation coefficients are Pearson linear partial coefficients after controlling for the common reference neurometabolite tCr; VC and PCC denote visual and posterior cingulate cortices; all neurometabolites are referenced to tCr and tissue-corrected; single and triple asterisks indicate $P < .05$ and $P < .001$ , respectively.*

### High-resolution temporal dynamics of GABA and Glx

To assess high temporal resolution dynamics of neurometabolites, we next combined transients across subjects, rather than across time. In visual cortex, we found change in GABA+, relative from the first measurement, was primarily negative and ranged from ±15% from the average concentration, while change in Glx was primarily positive and ranged from ±10% (**Fig. 4a**). These results are consistent with those from the previous analysis, except the higher temporal resolution obtained with the current approach revealed considerably larger changes in neurometabolite concentration than the previous estimates, which were likely obscured by smoothing measurements across the temporal window. In posterior cingulate cortex, we found that change in both GABA+ and Glx was primarily negative, and similar in amplitude (**Fig. 4b**).

**Figure 4.**
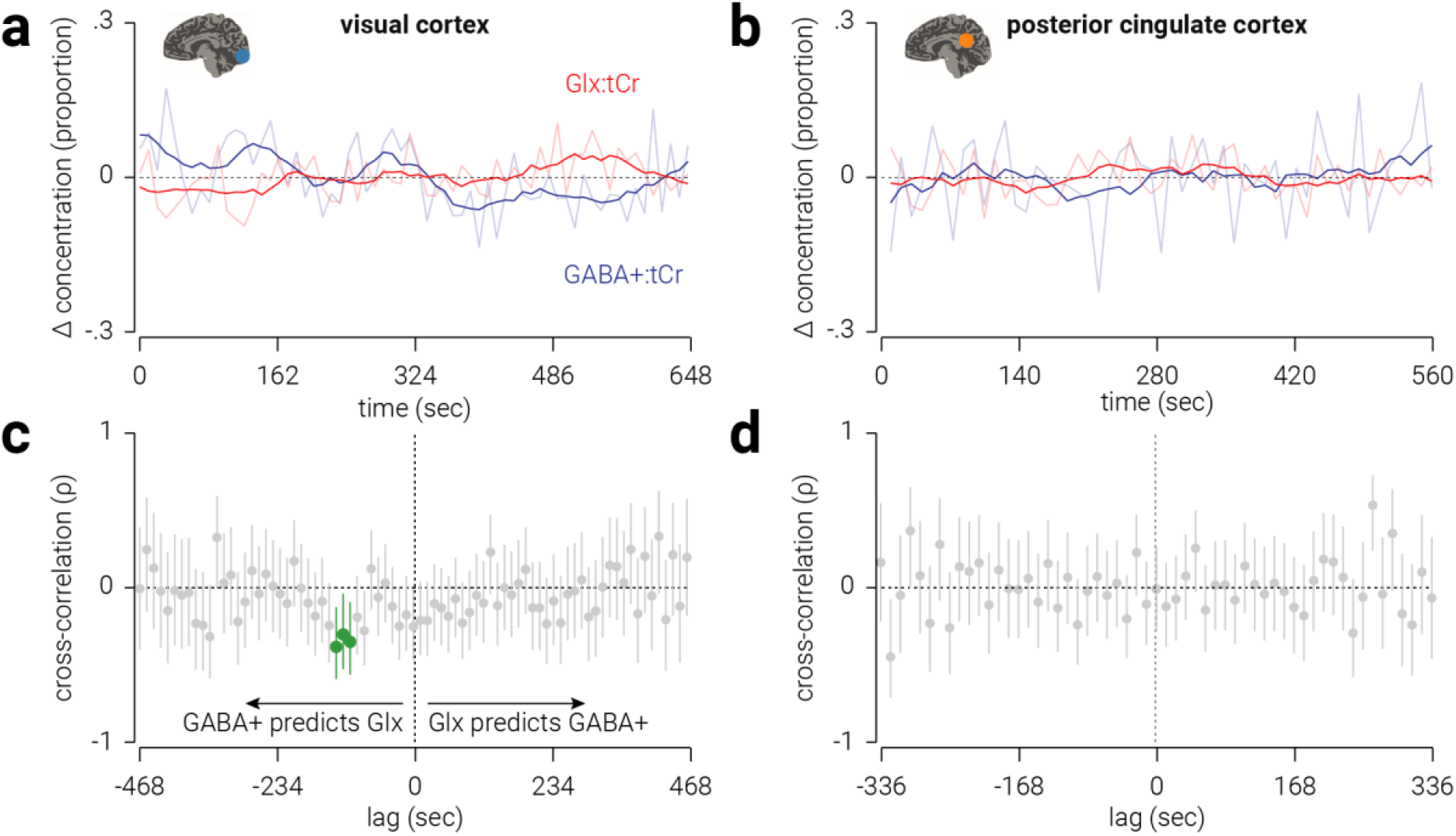
High-resolution temporal dynamics of neurometabolites in visual and posterior cingulate cortices. Change in GABA+ and Glx concentration, proportional to the average, measured from voxels targeting (**a**) visual and (**b**) posterior cingulate cortices. Semi-transparent lines indicate raw concentration estimates; for illustrational purposes, opaque lines indicate data smoothed using a window of eight time points. (**c-d**) Cross-correlations between GABA+ and Glx concentration measured from voxels targeting (**c**) visual and (**d**) posterior cingulate cortices. Lag values indicate the duration between when the GABA+ measurements were acquired and the Glx measurements were acquired. Correlations at negative lags indicate GABA+ concentration predicts Glx concentration and correlations at positive lags indicate Glx predicts GABA+. Vertical lines indicate 95% confidence intervals; cluster-corrected correlations that are significantly different than zero are highlighted in green. All values are referenced to tCr.

While GABA and Glx are thought to support opposing mechanisms in the central nervous system, i.e., inhibition and excitation, it seems reasonable to expect interactions between these neurometabolites. For example, Gln is a primary source of GABA synthesis (Patel et al., 2001; Paulsen et al., 1988; Rae et al., 2003). Indeed, we found that the static concentration of GABA+ and Glx were positively related in posterior cingulate cortex. However, we found no evidence for a relationship between the overall changes in these neurometabolites in either the visual or posterior cingulate cortices (**Table 2**). One reason for this may be that the relationship between these neurometabolites may only be observed at a high temporal resolution, but not averaged when across a 6 min period. To test this hypothesis, we used the high-resolution neurometabolite measurements to perform cross-correlation analyses on GABA+ and Glx concentration.

For visual cortex, we found that the concentration of GABA+ predicted that of Glx between 108-132 sec later (n(time points)=[55,54,53], Pearson *r*=[-.36,-.31,-.39], *P*=[.008,.024,.004]; **Fig. 4c**). This relationship was negative, that is, a positive/negative change in GABA+ predicted a later change in Glx in the opposite direction. By contrast, we found no periods of latency in which Glx predicted the concentration of GABA+. For posterior cingulate cortex, we found no periods of latency in which there was a significant relationship between GABA+ and Glx (**Fig. 4d**).

A possible concern is that the relationship between GABA+ and Glx was the influenced by their common reference neurometabolite (tCr). In particular, as both GABA+ and Glx were referenced to tCr, the relationship between them could be explained by an autocorrelation in the concentration tCr. However, we found the same pattern of results when tNAA, rather than tCr, was used as a reference neurometabolite (**Fig. 5a**). Another possible concern with this new analysis is that the signal-to-noise ratio is insufficient to yield valid measurements, e.g., due to reduced neurometabolite quantification accuracy. To assess the validity of the measurements, we compared the average FWHM and fit error of peaks produced in the high-resolution analysis with those produce in the low-resolution analysis. For visual cortex, we found no evidence for a difference in FWHM for any of the target neurometabolites (GABA+, *z*=-0.06, *P*=.948; Glx, *t*_(119)_=-5.08, *P*=1.4e_−6_; tCr, *z*=-1.10, *P*=.270). By comparison, fit error was lower for all target neurometabolites (GABA+, *z*=2.38, *P*=.017; Glx, *z*=4.21, *P*=2.5e_−5_; tCr *t*_(119)_=2.24, *P*=.027) modelled in the high-resolution analysis. Similarly, for posterior cingulate cortex, we found no evidence for a difference in either FWHM (GABA+, *z*=1.62, *P*=.105; Glx, *t*_(119)_=1.76, *P*=.080; tCr, *z*=1.71, *P*=.087); however, while we found no evidence for a difference in fit error for GABA (*z*=1.32, *P*=.187) and Glx (*z*=-0.55, *P*=.581), fit error was higher for tCr (*z*=-2.34, *P*=.019) in the high-resolution analysis. With the exception of fit error of the tCr peak in posterior cingulate cortex, these results support the validity of the signal strength and quantification accuracy in the high-resolution analysis.

**Figure 5.**
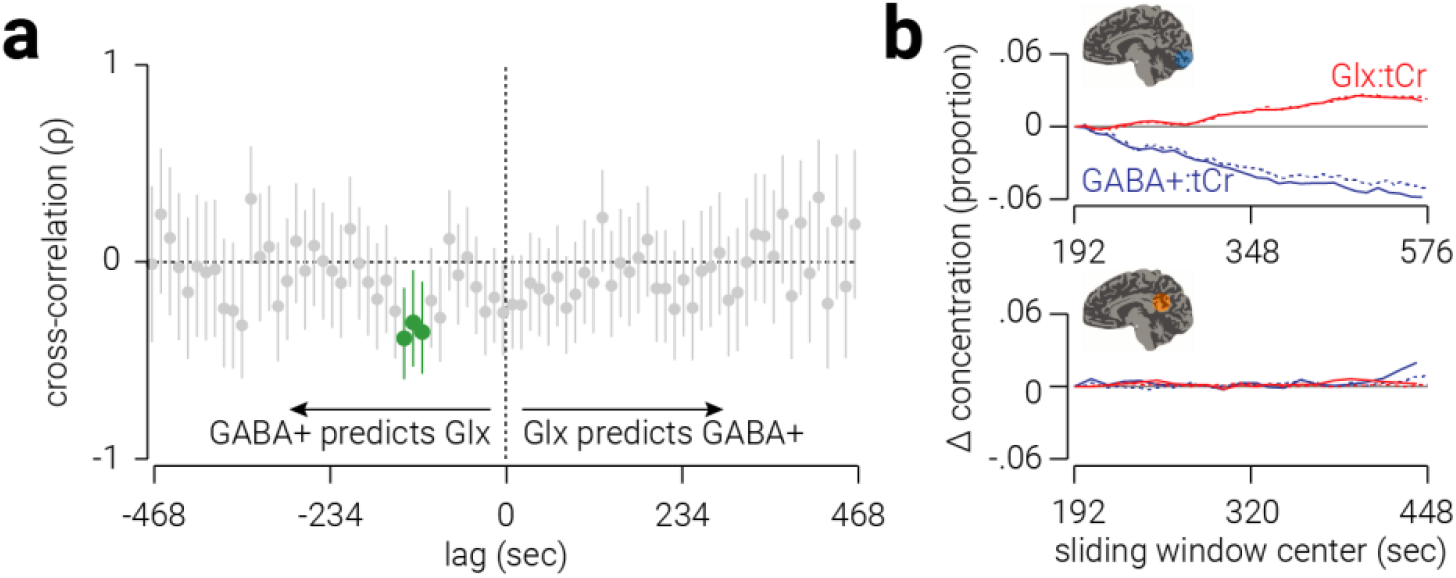
Controls for high-resolution analyses. (**a**) Cross-correlations between GABA+ and Glx, referenced to tNAA, measured from voxels targeting visual cortex. Vertical lines indicate 95% confidence intervals; cluster-corrected correlations that are significantly different than zero are highlighted in green, correlations at negative lags indicate GABA+ predicts Glx and correlations at positive lags indicate Glx predicts GABA+. (**b**) Comparison between results from sliding window method used in the low temporal resolution analysis (dashed lines) and sliding window applied to results from high-resolution analysis (solid lines) for (**b**, top) visual and (**b**, bottom) posterior cingulate cortices.

To further assess the validity of the measurements, we attempted to reproduce the results from the low-resolution analysis by applying a sliding window to the high-resolution neurometabolite trace. If the high-resolution measurements are valid, we would expect to find correspondence between the average results from the low-resolution analysis and those produced by applying a sliding window to the high-resolution measurements. For visual cortex, we found a high correspondence between the measurements produced by the two analyses for both GABA+ (n=33, Pearson *r*=.995, *P*=3.9e_−33_; Fig. 5b, top) and Glx (n=33, Pearson *r*=.995, *P*=5.5e_−32_). These results further validate the results of the high-resolution analysis in visual cortex. For posterior cingulate cortex, we found a correspondence between GABA+ measurements (n=22, Pearson *r*=.655, *P*=9.3e_−4_; Fig. 5b, bottom), but not Glx measurements (n=22, Pearson *r*=.182, *P*=0.418). The weaker correspondence for data from posterior cingulate cortex may indicate that the measurements produced by the high-resolution analysis in this region are less reliable. However, it is also likely because there was less variability in neurometabolite concentration in the low-resolution analysis of posterior cingulate cortex; thus, unlike in visual cortex, there may be insufficient variability to cross validate between the two measurements.

## DISCUSSION

MRS can be used in vivo to quantify neurometabolite concentration and provide evidence for the involvement of different neurotransmitter systems, e.g., inhibitory and excitatory, in sensory and cognitive processes. In MRS studies, temporal resolution is typically sacrificed to achieve sufficient a signal-to-noise ratio to produce a reliable estimate of neurometabolite concentration. Here we use novel analyses with large datasets to reveal the dynamics of GABA+ and Glx in visual and posterior cingulate cortices. We use a sliding window approach to show that when participants are at rest with their eyes closed, the concentration of GABA+ and Glx in visual cortex drifts in opposite directions, that is, GABA+ decreases while Glx increases over time. We then use a new method of combining MRS measurements across subjects, as opposed to time, to produce a high temporal resolution index of neurometabolite concentration. Using this approach, we find that in visual cortex, a change in the concentration of GABA+ predicts the opposite change in Glx ~120 sec later, e.g., an increase in GABA+ predicts a later reduction in Glx.

### Dynamic response of GABA and Glx in visual cortex

Several studies have investigated GABA and/or Glx/Glu concentration in visual cortex in response to different viewing conditions. Mekle et al. (2017) found a ~5% reduction in GABA concentration in response to visual stimulation. By contrast, while Kurcyus et al. (2018) reported that GABA was 16% lower when participants had their eyes open with no visual stimulation compared to when closed, they, like others (Bednařík et al., 2018, 2015; Mangia et al., 2007; Schaller et al., 2013), found no evidence for a difference in GABA in response to visual stimulation. Based on these somewhat inconsistent findings, one may infer that visual stimulation, or merely having the eyes open, leads to a reduction in the concentration of GABA in visual cortex. Extending this rationale, one could predict that closing the eyes should produce an increase in GABA. By contrast, we found the opposite result: during a 13 min period of resting in which participants’ eyes were closed, the concentration of GABA+ in visual cortex reduced on average by 5%. These results are consistent with previous work showing that monocular deprivation leads to reduced GABA concentration (~8%) in visual cortex, but not posterior cingulate cortex, relative a pre-deprivation baseline measurement (Lunghi et al., 2015). Thus, our finding that GABA is reduced when both eyes are closed may indicate that visual deprivation, either monocular or binocular, evokes a reduction in the concentration of GABA in visual cortex.

Previous observations of Glu concentration in visual cortex have been more consistent; several studies have shown Glx/Glu concentration increases (2-4%) in response to visual stimulation (Bednařík et al., 2018, 2015; Ip et al., 2017; Kurcyus et al., 2018; Lin et al., 2012; Mangia et al., 2007; Schaller et al., 2013). By contrast, here we found Glx increased when participants’ eyes were closed. Increased Glu in visual cortex, evoked by visual stimulation, has been linked to increased blood-oxygenation level dependent responses (Ip et al., 2017). Here we measured Glx, a complex comprising Glu and Gln, and previous 7T MRS work suggests that visual stimulation evoked changes in Glu, but not Gln, in visual cortex (Bednařík et al., 2018, 2015; Schaller et al., 2013). It is possible that the increase in Glx we found here, which occurred in the absence of visual stimulation, was driven by an alternative mechanism, one that is unrelated to BOLD activity and/or reflects changes in Gln rather than Glu. More work is needed it disambiguate changes in neurometabolite concentration that occur in visual cortex at different time scales and under different viewing conditions. For instance, future work could combine fMRI and MRS measurements to test whether the phenomenon observed here is related to BOLD activity (Ip et al., 2017).

Although the average concentration of GABA+ and Glx drifts in opposing directions, there were some participants who showed the opposite change in neurometabolite drift. One possible explanation for this is that the concentration of these neurometabolites oscillates at a relatively slow frequency (e.g., wavelength = 15 min), and entering a state of rest (with eyes closed) introduces the linear trend of neurometabolite concentration change indicated by the average, which is summed with the larger oscillatory changes. If such oscillatory behaviour was occurring, it would be unlikely to be detected in the high-resolution analysis, as troughs/peaks would be obscured by phase differences between participants.

Continuous unidirectional neurometabolite drift is not sustainable, i.e., neurometabolite concentration must maintain some degree of homeostasis. Thus, it seems likely that the change in concentration induced by resting only continues for a fixed period of time, before stabilising and possibly returning to ‘baseline’ levels. Indeed, this appeared to occur for Glx, where there is no additional increase after ~500 s, and the decrease is GABA+ appears to lessen after 400 s. Future work is needed to further understand dynamic neurometabolite activity over longer periods of time.

To produce a high temporal resolution measure of neurometabolite concentrations, we applied a novel approach in which we combined MRS transients across subjects, rather than time. This approach yields a single measurement of neurometabolite concentration as a function of time; thus, we cannot test the statistical significance of the changes observed. However, it is reasonable to assume that the changes estimated using sliding window or static approaches underestimate the true magnitude of change, due to averaging. The measurements produced here by combining transients across subjects provide an indication of the true magnitude of change in GABA+ and Glx during the scan: up to 30% for GABA+ and 20% for Glx.

### Dynamic relationship between GABA and Glx

A common finding among studies of GABA and Glu in visual cortex is that these neurometabolites change in opposite directions, and not in the same direction (Kurcyus et al., 2018; Mekle et al., 2017; Rideaux et al., 2019). This pattern would suggest an interdependency between the two neurometabolites. Given that Gln is a primary source of GABA synthesis (Patel et al., 2001; Paulsen et al., 1988; Rae et al., 2003), one may expect that a change in the concentration of GABA may result in a corresponding change in Glx, or vice versa. In line with this, our high temporal resolution analysis of the data revealed a striking cross-correlation between these neurometabolites in visual cortex. Specifically, we found that the concentration of GABA+ predicts the concentration of Glx ~120 sec later. This relationship, which accounts for up to ~20% of the variance of Glx, is obscured by conventional approaches of MRS analysis; indeed, we found no evidence for a relationship between the overall change in GABA+ and Glx.

A limitation of MRS is that it measures the total concentration of neurometabolites within a localized region and cannot distinguish between intracellular and extracellular pools of GABA. It is generally thought that intracellular vesicular GABA drives neurotransmission (Belelli et al., 2009), whereas extracellular GABA maintains tonic cortical inhibition (Martin and Rimvall, 1993). Synthesis of intracellular GABA from Gln via the GABA-Gln cycle occurs on the scale of milliseconds, so it seems unlikely that this metabolic association could explain the predictive relationship between GABA and Glx concentration 120 sec later. Instead, this relationship may reflect changes in the level of extracellular GABA, followed by relatively sluggish changes in the concentration of Glu to maintain the balance of inhibition and excitation.

### Clinical relevance

An imbalance between excitation and inhibition, in particular between Glu and GABA, is thought to underlie a range of neurological and psychiatric disorders including epilepsy (Bradford, 1995; Olsen and Avoli, 1997), autism (Chao et al., 2010; Markram and Markram, 2010; Rubenstein and Merzenich, 2003; Vattikuti and Chow, 2010), and schizophrenia (Kehrer et al., 2008). Here we demonstrated that it is possible to use MRS to measure changes in neurometabolite concentration at relatively high temporal resolution, and in the process of doing so, we revealed a novel predictive relationship between GABA+ and Glx in visual cortex of healthy human participants. More broadly, this method could be used to reveal other trends and relationships between neurometabolites that have thus far been obscured by the classic ‘static’ measurement approach. Uncovering the dynamics of the excitation-inhibition balance in this way may be instrumental in understanding unhealthy brain function. However, a caveat to this approach is that it requires a relatively large amount of MRS data. Here, we used a large number of subjects to meet this requirement; however, this may not be appropriate for clinical studies in which suitable subjects are more difficult to recruit. In this case, a modified event-related version of the method used here may be effective (Branzoli et al., 2013). That is, events (e.g., a 2 min stimulus presentation) could be repeated multiple times over a long period, separated by baseline resting intervals. Transients could then be aligned according to the time course of the event and averaged together. This would provide the same signal-to-noise improvement reached in the current study, while requiring fewer participants.

### Conclusion

The relatively low signal-to-noise of MRS measurements has shaped the types of questions that the technique has been used to address. Here we overcome this limitation by combining data from large cohorts to examine the dynamics of GABA and Glx concentration in visual cortex. Through use of existing and novel analyses, we reveal opposing dynamic shifts in GABA and Glx in visual cortex while participants are at rest. Further, we demonstrate a predictive relationship between GABA and Glx in visual cortex. This study exposes temporal trends of primary neurotransmitters in visual cortex, and more generally, these findings demonstrate the feasibility of using MRS to investigate in vivo dynamic changes of neurometabolites.

## Acknowledgements

The work was supported by the Leverhulme Trust (ECF-2017-573) and the Issac Newton Trust (17.08(o)).

